# Agile manufacturing of PDMS microfluidic devices for cell culture without a clean room

**DOI:** 10.64898/2026.11.07.765050

**Authors:** Jean Abel Merino, Cristian Angeli, Juan Manuel García-Arcos

## Abstract

Microfluidic devices control fluid flow and the cellular microenvironment, and their fabrication usually relies on photolithography in a clean room. For devices with features of hundreds of micrometers to millimeters, this route is slow and costly, especially during prototyping, when the design changes after each test. Here, we describe and benchmark a workflow to fabricate polydimethylsiloxane (PDMS) devices from laser-cut master molds, with equipment accessible to most biology laboratories. We cut three inexpensive materials, poly(methyl methacrylate) (PMMA), polystyrene and polyvinyl chloride (PVC) tape, over a grid of laser power and speed and measured each cut by scanning electron microscopy. Cut depth and width increased linearly with the laser energy delivered per unit length, which gives a simple rule to choose the settings for each material. We then bonded the PDMS replicas to glass without a plasma cleaner, by stopping curing early, once the replica could be demolded, and completing it in contact with the glass. Without any surface treatment, these devices held up to about 400 mbar; activating the glass with a handheld corona device raised this to about 900 mbar, enough for perfusion. The master material set the surface of the device: PMMA gave smooth replicas, whereas PVC tape gave rough walls. A three-channel device molded on PMMA held two cell-laden gels in place with phase guides and supported the co-culture of cancer cells and fibroblasts under perfusion. Further analysis of flow laminarity and different mixer designs confirmed the robustness of this method. This workflow takes a digital design to a working device within a day and lets biology laboratories adapt microfluidic devices to their experiments without the need of a clean room.

## Introduction

Microfluidic devices control small volumes of fluid and are established tools in chemistry, biophysics and cell biology [1]. At these length scales, flow is laminar, and transport is set by channel geometry, diffusion and the applied flow. Microfluidic devices can therefore expose cells to defined chemical environments, confinement, shear or time-varying stimuli [2]. Polydimethylsiloxane (PDMS) remains one of the most widely used materials for academic devices because it is transparent, elastomeric, permeable to gases and easy to mold [3].

PDMS devices are usually made by soft lithography: a master mold is patterned in photoresist, typically SU-8 on a silicon wafer, and PDMS is cast against it [4,5]. Soft lithography is the method of choice for features of a few micrometers, but it requires a photomask, a clean room and trained users. These requirements are a barrier for biology laboratories that use microfluidics as one tool among many, and they slow down development, when the design changes after each test [6–9].

Many biological experiments need channels and chambers of several hundred micrometers or more. At these dimensions, photolithography offers more resolution than needed, and each new design repeats mask design, resist processing, exposure and development. Alternatives include laser machining, cutting of adhesive films (xurography), micromilling and 3D printing [7,8,10–20], and photolithography with UV-curable nail gel [21]. All aim to shorten the path from a modified digital design to a new device.

Laser cutters suit this aim because they translate a vector drawing directly into a polymer structure. Laser-patterned tape has served as a master for PDMS replica molding [10], including molds for multilayer devices with valves [11,12], and laser-machined poly(methyl methacrylate) (PMMA) has been used both as the device and as the mold [7,13,14,22]. Laser cutting, however, is a thermal process: the geometry of a cut depends on laser power, translation speed, focus and the polymer [23–25], and cuts can have tapered walls, melted edges and rough surfaces [7,13,14]. The dimensions of a design therefore do not transfer directly to the master. A practical method needs a calibration that links the laser settings to the dimensions obtained in each material, and a test of how the surfaces left by the laser affect the device.

The second bottleneck is bonding. Oxygen plasma activates PDMS and glass, which then form covalent siloxane bonds on contact [5,26]. Plasma cleaners, however, are not standard equipment in biology laboratories. Handheld corona devices are a cheaper source of surface activation [27], and partially cured PDMS layers bond to each other without activation [28], because their network can still cross-link across the interface. This suggests that curing time could replace the plasma cleaner: PDMS cured just enough to be demolded might bond to glass if its curing is completed in contact with it.

Here, we describe a workflow to fabricate PDMS devices from laser-cut masters, and we benchmark the steps that decide whether a device works. We mapped cut width and depth in three inexpensive master materials over a grid of laser power and speed, and found that the energy delivered per unit length predicts the cut. We then tested whether partially cured PDMS bonds to glass-bottom dishes without a plasma cleaner, either without surface treatment or after corona activation of the glass. Finally, we compared the surfaces left by rigid sheets and by tape, and tested how they affect three devices: a three-channel device for co-culture in gels, a replica of a commercial channel slide, and a two-inlet device that mixes or switches two solutions. The workflow suits features of hundreds of micrometers or more, when iteration speed matters more than micrometer resolution.

## Results

### A workflow to fabricate PDMS devices from laser-cut masters

We built the workflow around equipment that a biology laboratory can own: a benchtop CO_2_ laser cutter, an oven and a hotplate (Fig 1). The master, the mold on which PDMS is cast, is cut from commercial polymer sheets or tape, and the PDMS steps are the same for all materials. For rigid sheets, a double-sided adhesive film is laminated to the sheet, the design is cut through, and the released features are attached to glass (Fig 1A). For tape, the tape is applied to glass, cut through and the excess peeled off, which avoids the transfer step. PDMS is then cast on the master, cured, peeled, punched and bonded to a glass-bottom dish (Fig 1B). A master is changed by editing the vector design and cutting again, without a photomask.

**Fig 1.**
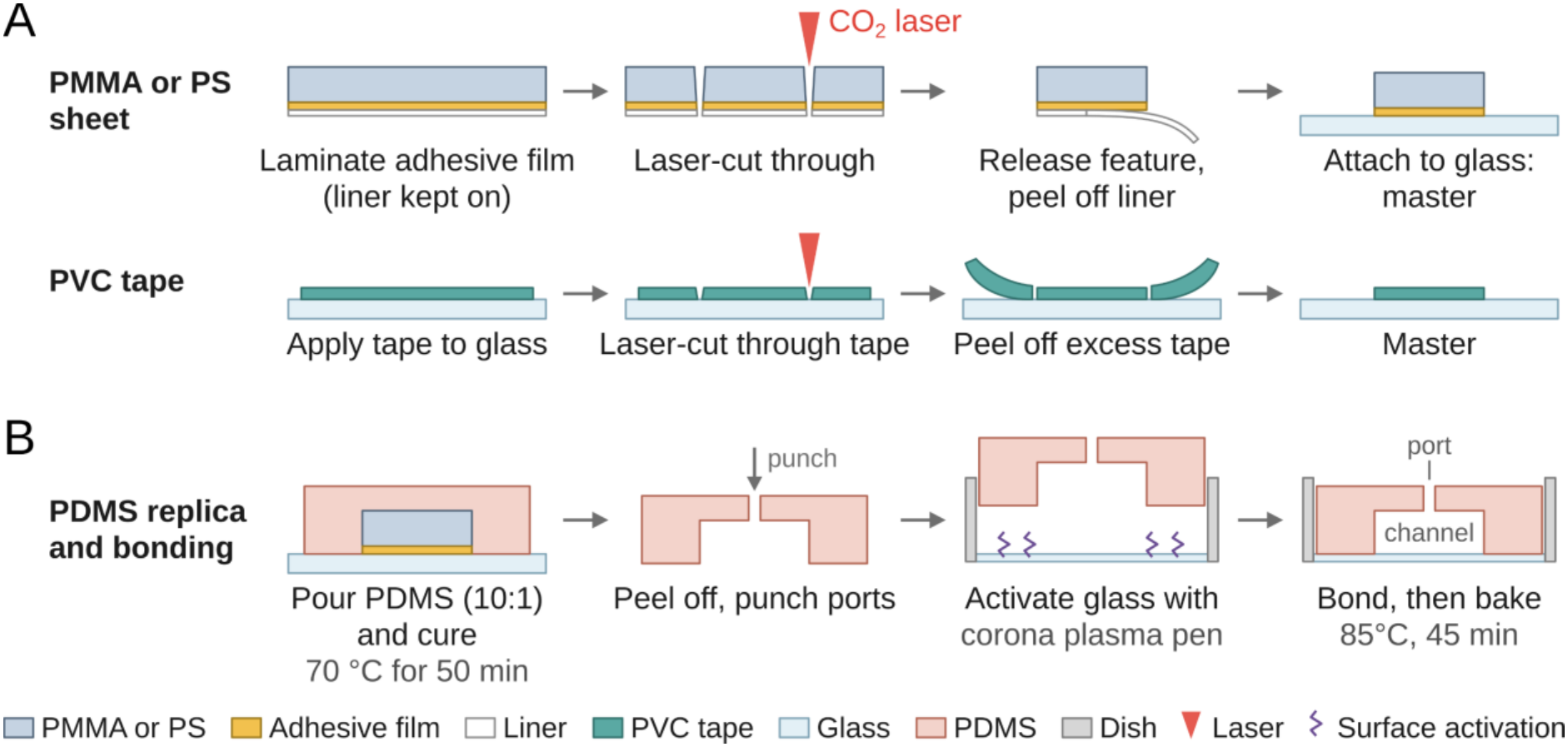
Fabrication of PDMS devices from laser-cut master molds. (A) Master fabrication. Rigid sheets (PMMA or PS): a double-sided adhesive film is laminated to the sheet, keeping the liner on its outer face; the design is cut through the sheet with a CO_2_ laser; the features are released, the liner is removed, and the features are attached to a glass support. PVC tape: the tape is applied to the glass support, the design is cut through the tape, and the excess tape is peeled off. (B) PDMS replication and bonding. PDMS is poured over the master and partially cured (70 °C, 50 min); the replica is peeled off, and ports are punched; the glass of the dish is activated with a handheld corona device (optional); the replica is placed on the dish and bonded on a hotplate (85 °C, 45 min). Not to scale.

We compared three materials. PMMA is rigid, widely available and well characterized for laser machining [7,14]. Polystyrene (PS) is inexpensive and is used for thermoplastic microdevices [17] and culture dishes, but it responds differently to the laser [29] and has rarely been tested as a master material. PVC tape is cut directly on its glass support [10]. For any of these materials, the master is predictable only if the effect of the laser settings on the cut is known, so we measured it first.

### Line energy sets the width and depth of laser cuts

Two dimensions decide whether a laser setting gives a usable master: the depth H, which must reach through the material to release a feature, and the width W, the material removed around the designed contour. We cut a test pattern in 1-mm PMMA, 1-mm PS and 0.15-mm PVC tape at 49 combinations of power and speed per material and measured H and W on scanning electron microscopy (SEM) cross-sections (Fig 2; S1–S3 Fig). The tape, much thinner than the standard sheets, was cut at lower power and higher speed.

**Fig 2.**
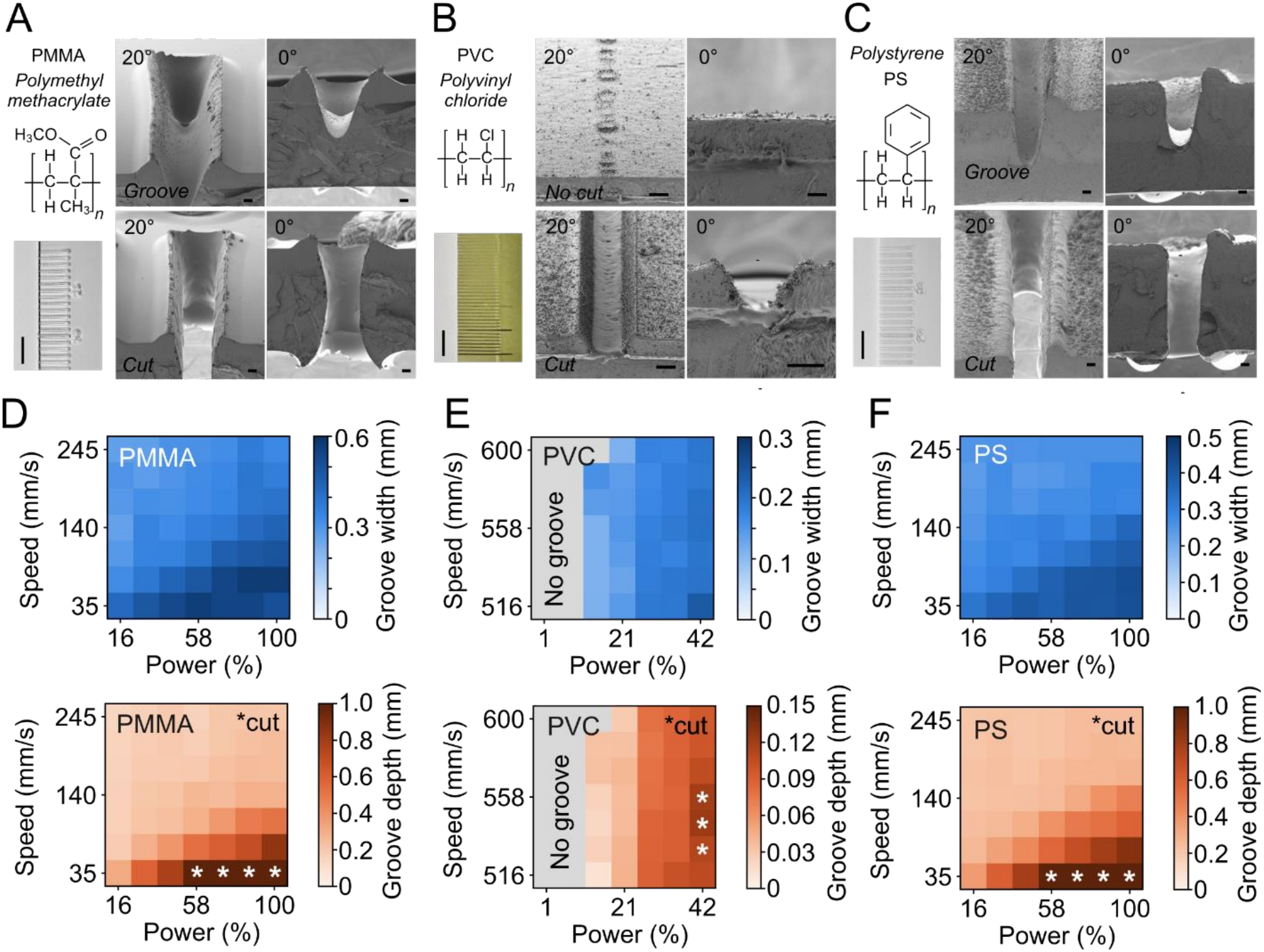
Laser power and speed set the width and depth of cuts in PMMA, polystyrene and PVC tape. (A, B, C) Left, chemical structure and photograph of the test cuts<u>, s</u>cale bars, 6 mm for PMMA (A), PVC (B) and PS (C). Right, SEM images of a groove (no cut in PVC tape) and of a cut through the material, at 20° tilt and in cross-section (0°) for PMMA (A), PVC (B) and PS (C), scale bars, 100 µm. PMMA and PS groove, 100% power and 105 mm/s; PMMA and PS cut, 100% and 35 mm/s; PVC tape, no cut at 1% and 600 mm/s and cut at 42% and 516 mm/s. (D, E, F) Cut width W (blue) and depth H (orange) for each combination of laser power (% of 55 W) and translation speed; one cut per condition. Asterisks, cuts through the 1-mm sheet or through the 110-µm PVC film; gray, no measurable groove. All cross-sections in S1–S3 Fig. Data in S1 Data.

Each material had its own cutting window (Fig 2D–2F). In both 1-mm sheets, the laser cut through only at the lowest speed and at 58% power or more; at all other settings it left a groove. In tape, low power left no measurable groove, and the laser cut through the PVC film only at the highest power. Within each material, depth and width increased linearly with the nominal line energy E = P/v, the energy delivered per unit length (S4A and S4B Fig), in agreement with earlier measurements and models of laser-machined channels in PMMA [7,23,24]. In the sheets, depth fell below this line at high power (S4C Fig), suggesting a stronger effect of power over speed. Power and speed can therefore be traded at approximately constant P/v, and one grid of cuts is enough to calibrate a material.

Through-cuts define the master. They were about 0.5 mm wide at the top in PMMA and 0.4 mm in PS, and narrower at the base, more so in PS (S4D Fig). PMMA largely depolymerizes and vaporizes under laser heating [30], whereas PS also melts [29], which may explain the difference. Through-cuts in tape were about 0.2 mm wide (Fig 2E). Because a feature loses half a cut width on each side, a master is narrower than its design by one cut width. In practice, the user chooses the lowest line energy that cuts through the material and widens the design by one cut width.

### Partially cured PDMS bonds to glass without a plasma cleaner

We next addressed bonding without a plasma cleaner. We reasoned that partially cured PDMS still carries unreacted cross-linking groups, which could bond to glass if curing is completed in contact with it. This hypothesis predicts a window of curing times: below it, the replica is too soft to be demolded; above it, the PDMS is too cross-linked to bond. We tested this prediction on a chip with one straight channel (Fig 3A). Replicas cured at 70 °C for 10 to 120 min were placed on glass-bottom dishes and heated on a hotplate to complete curing. We then raised the pressure in the channel in steps until the device leaked, a common test of bond strength [31].

**Fig 3.**
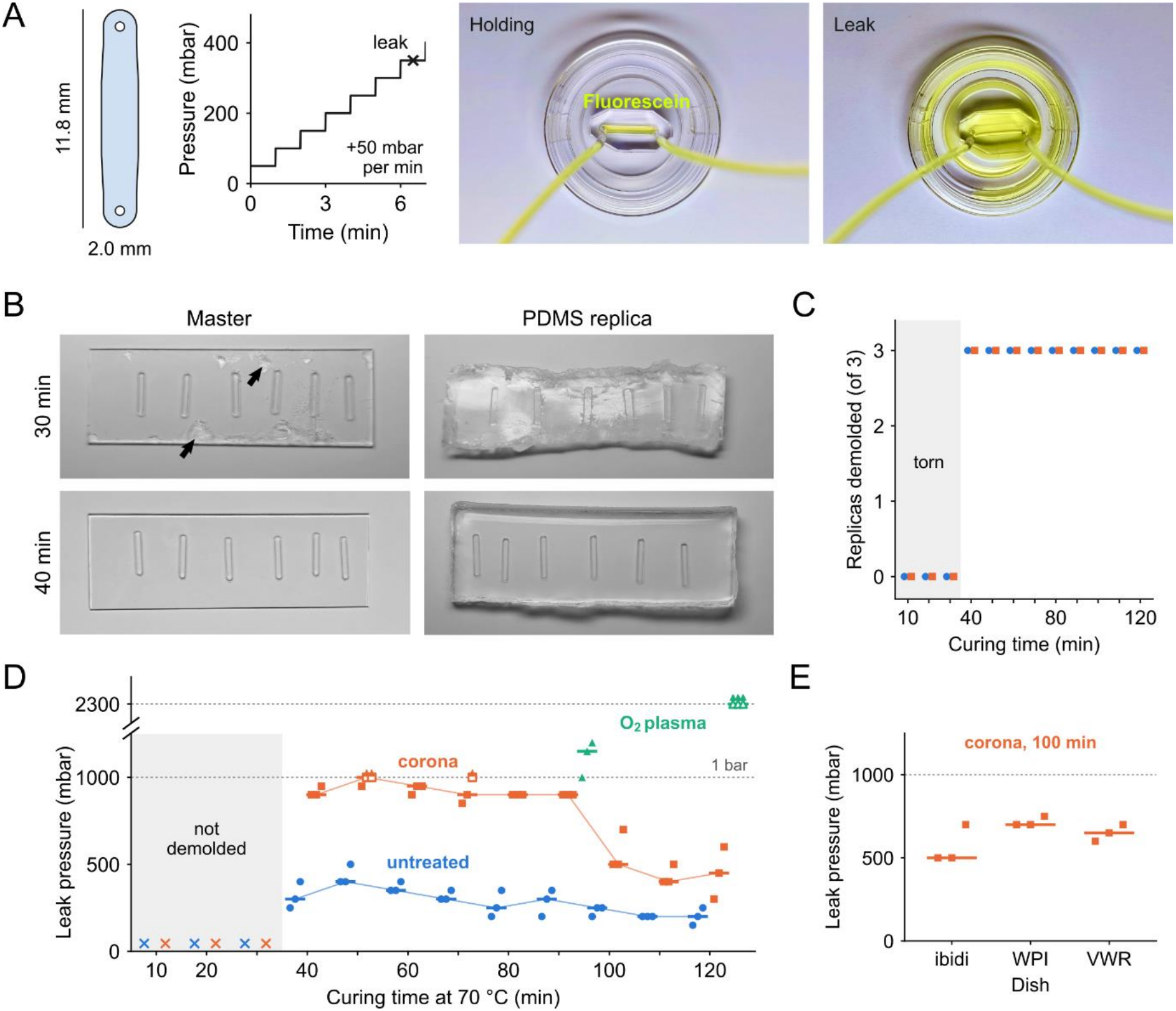
Partially cured PDMS bonds to glass, and corona activation strengthens the bond. (A) Test chip (straight channel, 11.8 × 2.0 mm in the design), pressure protocol (50-mbar steps every minute until the device leaks) and photographs of a device holding pressure and of a leak (fluorescein). (B) PMMA master and PDMS replica after 30 min of curing (replica torn, PDMS left on the master, arrows) and after 40 min (released intact). (C) Replicas demolded intact, of 3 per curing time, without treatment (blue) and with corona activation (orange); gray, curing times at which no replica could be demolded. (D) Leak pressure as a function of curing time at 70 °C for devices bonded on a hotplate to Ibidi glass-bottom dishes without treatment (blue circles) or after corona activation of the dish (orange squares), and for devices bonded after oxygen plasma (green triangles; tested with a 2.3-bar pressure generator, upper part of the broken axis). ×, not demolded; open symbols with arrowheads, devices that held the maximum pressure (dotted lines); lines connect the medians. Without treatment, leak pressure decreased with curing time (Spearman ρ = −0.76, P < 0.001). Corona-activated devices leaked at higher pressures than untreated devices (Mann–Whitney U test, P < 0.001, 40–120 min), and at lower pressures after 100–120 min than after 40–90 min of curing (Mann–Whitney U test, P < 0.001). (E) Leak pressure on three brands of 35-mm glass-bottom dish after 100 min of curing and corona activation (Krus-kal–Wallis test, P = 0.14). Points, devices (n = 3 per condition); bars, medians. Data in S1 Data.

The lower bound was sharp: replicas cured for 30 min or less tore during demolding, whereas all replicas cured for 40 min or more were released intact (Fig 3B and 3C). Without surface activation, these replicas bonded to the glass, and the bond weakened with longer curing, as predicted (Fig 3D). This bond needed no surface treatment, but held at most about 400 mbar.

For higher pressures, we activated the glass with a handheld corona device, about an order of magnitude cheaper than a plasma cleaner. Corona activation strengthened the bond at every curing time. Devices cured for 40 to 90 min leaked only at about 900 mbar, and longer curing weakened the bond, which marks the upper bound of the window (Fig 3D). Oxygen plasma, in contrast, bonded fully cured PDMS best, as expected in its standard use (Fig 3D). The bond was similar on three brands of glass-bottom dish, tested at the upper bound of the window, where it should be most sensitive to the substrate (Fig 3E). Partial curing therefore replaces the plasma cleaner: without treatment for devices that operate at low pressure, and with corona activation for devices that operate below about 900 mbar. For all following devices, we cured PDMS for 50 min, in the middle of the window, and bonded it after corona activation.

### The master material sets the surface of the device

Our calibration so far describes the cut in cross-section, but not the surfaces that the PDMS replicates. We compared these surfaces on masters of one design, cut in 0.5-mm PMMA and in three layers of PVC tape (Fig 4A and 4B). The design is a three-channel device in which two phase guides, ridges across the channel ceiling, pin the front of a liquid and separate the channels without a wall [32]. The phase guides were cut partway into the top of the master.

**Fig 4.**
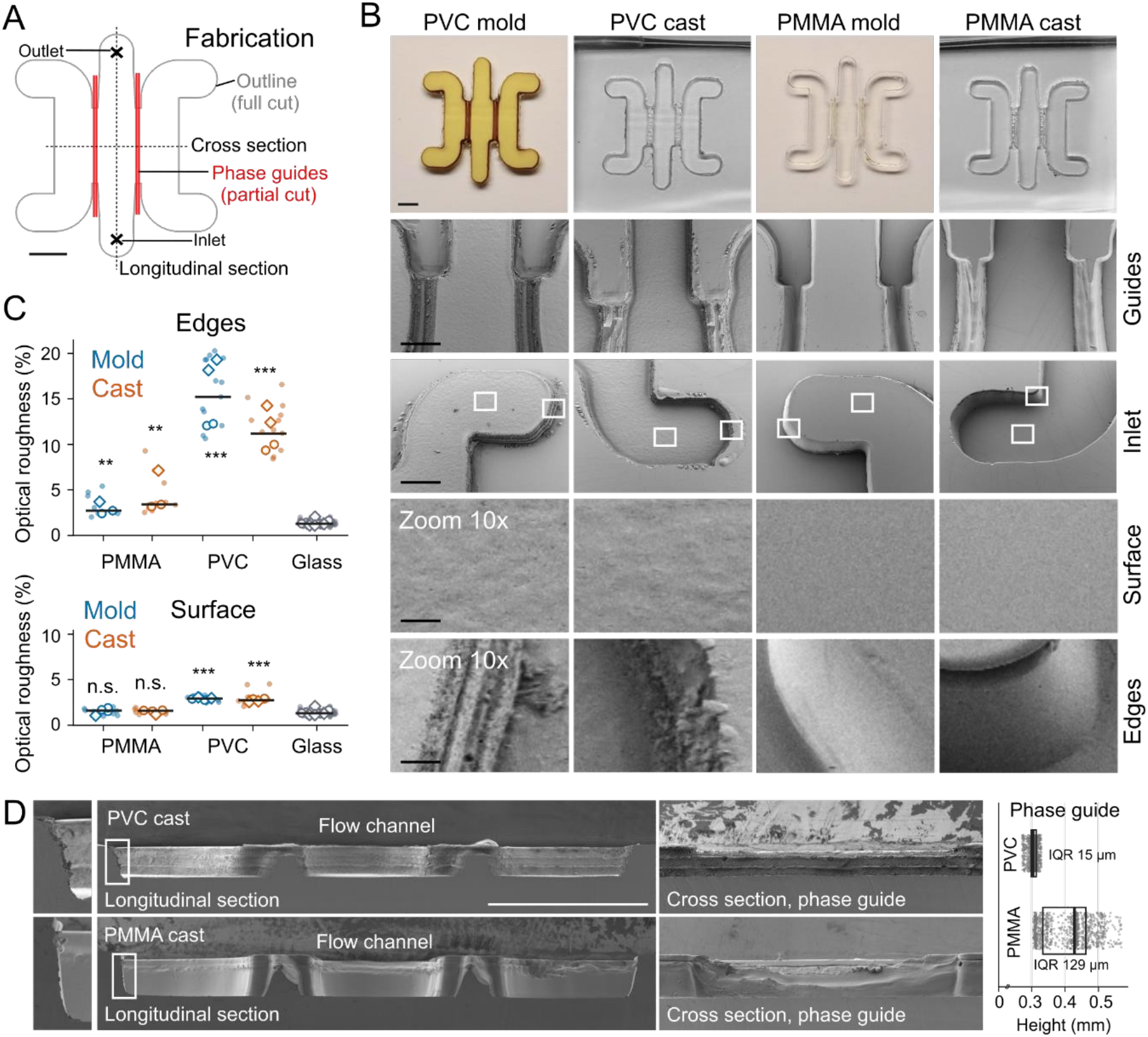
The master material sets the surface of the device. (A) Design of the three-channel device: outline cut through the material (gray) and phase guides cut partway (red); dotted lines, positions of the sections in D. (B) Masters (molds) in PVC tape and PMMA and their PDMS casts: photographs (top) and SEM images (20° tilt) of the phase guides, an inlet (white squares, regions enlarged 10× below), a flat surface and an edge. (C) Roughness of edges (top) and flat surfaces (bottom) of molds (blue) and casts (orange), and of glass (gray; glass support next to molds, PDMS molded against glass next to casts): relative variation of the SEM signal at scales of about 10–60 µm. Dots, regions; open symbols, image medians from two imaging sessions (circles, diamonds); lines, means of the image medians. Mann–Whitney U test of image medians against glass (3–4 images per condition, 16 for glass): n.s., P ≥ 0.05; **, P < 0.01; ***, P < 0.001. (D) Longitudinal sections of PVC and PMMA casts (white boxes, enlarged at left) and cross-sections through a phase guide; right, channel height under the phase guide along the longitudinal section (box, interquartile range; line, median). Scale bars, 1 mm (A, B and D). Data in S1 Data.

The two masters had different surfaces, and the casts reproduced each of them (Fig 4B). Flat PMMA surfaces were as smooth as glass and PMMA edges only slightly rougher, whereas tape was rough on both flat surfaces and edges (Fig 4C). In the casts, each phase guide formed a continuous ridge across the channel, with a regular height along the tape cast but not along the PMMA cast (Fig 4D). PMMA masters therefore give smoother devices, and tape masters more regular partial cuts due to the layering. We next tested whether these surfaces support the two functions that culture devices need: stable compartments and predictable flow.

### Phase guides hold cell-laden gels in place for perfused co-culture

We used the device molded on PMMA to culture two cell types in adjacent gels, separated by a perfused medium channel (Fig 5A). A549 lung adenocarcinoma cells expressing mCherry were loaded into one lateral channel and MRC-5 lung fibroblasts into the other, each in a gel of Matrigel and SiR-labeled collagen I. The phase guides stopped the front of each gel, so that each gel filled its own channel without entering the central one, and the two gels could be loaded one after the other (Fig 5B) [32,33]. After polymerization, the central channel was connected to a closed perfusion loop with a peristaltic pump inside the incubator (Fig 5C).

**Fig 5.**
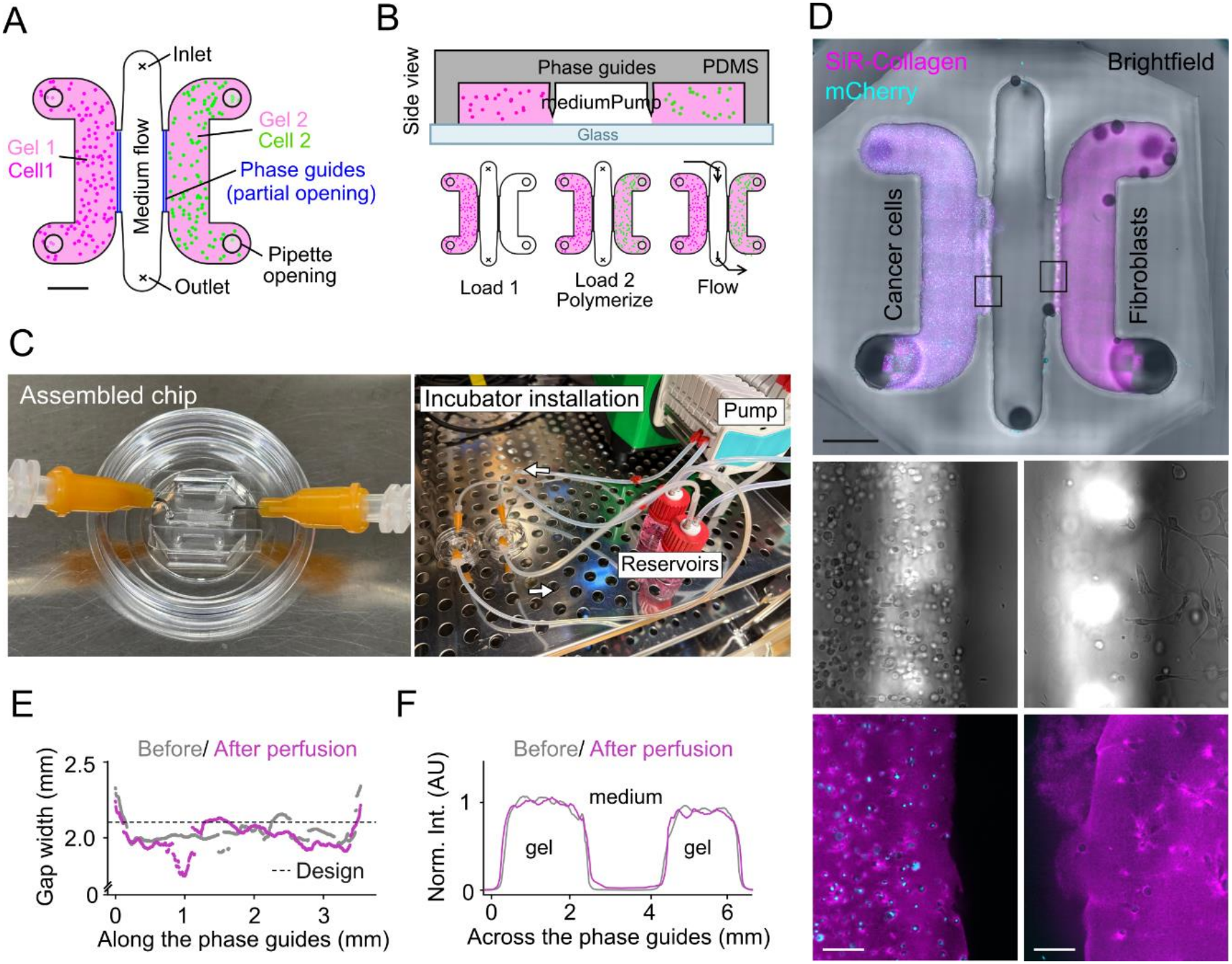
Phase guides hold two cell-laden gels in place for perfused co-culture of primary fibroblasts and cancer cells. (A) Device: two lateral channels, each holding a cell-laden gel (gel 1 with A549 cancer cells, gel 2 with MRC-5 fibroblasts), separated from the central medium channel by phase guides (blue). Gels are loaded through the pipette openings; medium flows from inlet to outlet. (B) Side view and loading sequence: gel 1 is loaded, then gel 2; both polymerize, and the central channel is perfused with medium. (C) Device bonded to a 35-mm glass-bottom dish with needles in the inlet and outlet (left), and closed perfusion loop in the incubator with the peristaltic pump and medium reservoirs (right; arrows, direction of flow). (D) Bottom view of the device: overlay of SiR-collagen (magenta), mCherry (cyan; cancer cells) and brightfield; black boxes, regions at the two phase guides enlarged below in brightfield (middle) and fluorescence (bottom). (E) Gap between the two gel fronts along the phase guides, before (gray) and 12h after the start of perfusion (magenta); dashed line, distance between the phase guides in the design (2.13 mm). (F) SiR-collagen intensity across the device before (gray) and 12h after the start of perfusion (magenta), normalized to 0 in the PDMS and 1 in the gel and averaged over the phase-guide section. One device. Scale bars, 1 mm (A and D, overview) and 100 µm (D, enlargements). Data in S1 Data.

Compartments are useful only if they stay separate under flow. The gap between the two gels was maintained after the start of perfusion (Fig 5E and 5F), and the two cell types remained in their own gels along the phase guides (Fig 5D). The device therefore supports perfused co-culture of two cell types in adjacent gels.

### The rough walls of tape masters do not disturb laminar flow

Tape masters had further defects. The edges of multilayer tape were layered and rough, the casts reproduced this texture, and the PDMS could fracture along it (S5A Fig). Adhesive left on a master that was not thoroughly cleaned with solvent was also reproduced in the cast (S5B and S5C Fig). Rough walls could, in addition, perturb the flow near the walls, which matters for experiments that rely on defined shear or transport. We therefore tested the flow in a channel molded on tape.

We replicated one channel of a commercial slide (ibidi µ-Slide VI 0.4) with a master of three layers of PVC tape and compared the flow in the replica and in the original by particle image velocimetry (PIV) [34] (Fig 6A and 6B). Both channels had the flat velocity profile of a wide, shallow channel, and the velocity scaled with the flow rate (Fig 6C and 6D). Vorticity was confined to the side walls in both channels (Fig 6E and 6F), and neither the velocity fluctuations in the center nor the variation of vorticity along the walls was larger in the replica (Fig 6G and 6H). The rough tape walls therefore did not disturb laminar flow.

**Fig 6.**
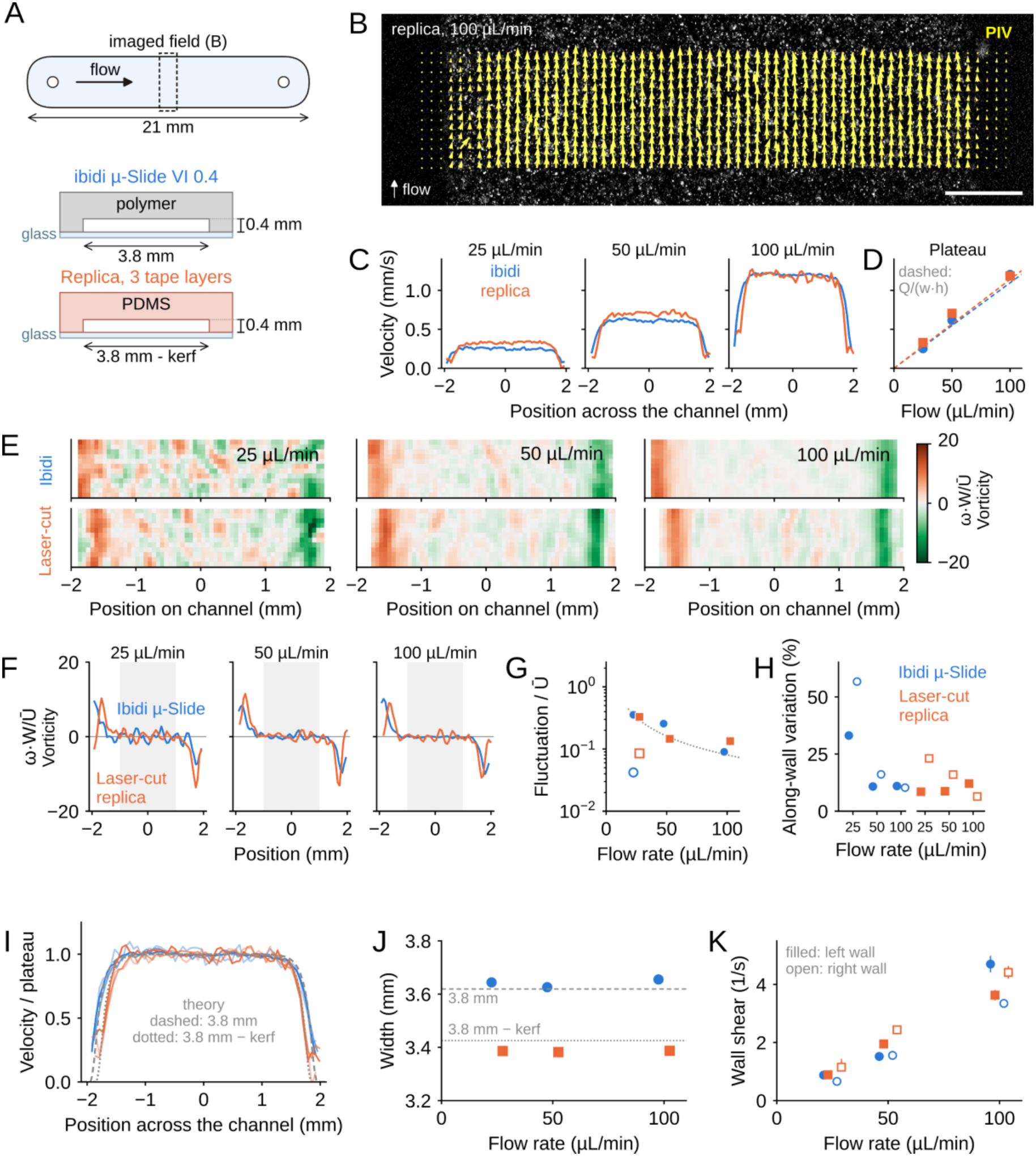
The rough walls of a tape-molded channel do not disturb laminar flow. (A) Top view of the channel (3.8 × 21 mm) with the imaged field, and cross-sections of the commercial slide (ibidi µ-Slide VI 0.4; polymer on glass, 0.4 mm high) and of the replica (PDMS on glass, molded on three layers of PVC tape). (B) Fluorescent 1-µm beads in the replica at 100 µL/min, with the mean PIV vectors (yellow); flow upward. Scale bar, 0.5 mm. (C) Streamwise velocity across the channel, averaged along the flow, at 25, 50 and 100 µL/min. (D) Plateau velocity (mean over the central 2 mm) as a function of flow rate; dashed lines, mean velocity expected from the flow rate and the nominal cross-section. (E) Time-averaged vorticity, normalized as ωW/Ū (W, channel width; Ū, mean velocity expected from the flow rate), in the commercial slide (top) and in the replica (bottom), on one color scale (green, negative; orange, positive). (F) Vorticity across the channel, averaged along the flow; gray band, central 2 mm. (G) Root-mean-square velocity fluctuation in the central 2 mm relative to Ū; dotted line, PIV noise; open symbols, after subtraction of the noise (25 µL/min). (H) Variation of the time-averaged wall vorticity along the flow at the left (filled) and right (open) walls. (I) Velocity normalized to the plateau at the three flow rates (darker, higher flow rate), with the exact solution for laminar flow in a rectangular channel of the design width (3.8 mm, dashed) and of the design width minus one cut width (dotted). (J) Width of the profiles at half the plateau velocity; lines, the same two solutions. (K) Shear rate at the left (filled) and right (open) walls, from the slope of the velocity profile; bars, 95% confidence intervals from resampling of the PIV fields. Blue circles, commercial slide; orange squares, replica. One recording per channel and flow rate (34–52 PIV fields each). Data in S1 Data.

The replica also allowed us to measure the width of the groove, as this causes a change in the PDMS dimensions compared to the nominal design. The flow field was narrower than designed, by about one cut width in tape (Fig 6I and 6J; Fig 2E). The same flow rate therefore gives a slightly higher shear stress on the floor of the replica, whereas the shear rates at the side walls were similar in both channels (Fig 6K). A tape replica thus reproduces the flow of a commercial channel, provided that the design is widened by one cut width, known as kerf.

### Laser-cut mixers mix or switch two solutions

Because laser-cut walls left laminar flow unchanged, we then built a device whose function depends on flow through several elements in series: two inlets, a mixer, a flow equalizer and a culture chamber (Fig 7A). Such devices deliver a defined mixture of two solutions to cells, or change it over time [35]. Because a new master is cut in minutes, we made two versions in PMMA, with a serpentine mixer or with a staggered herringbone mixer, whose grooves in the channel ceiling fold the two streams [36] (Fig 7B). One inlet carried fluorescein and the other water.

**Fig 7.**
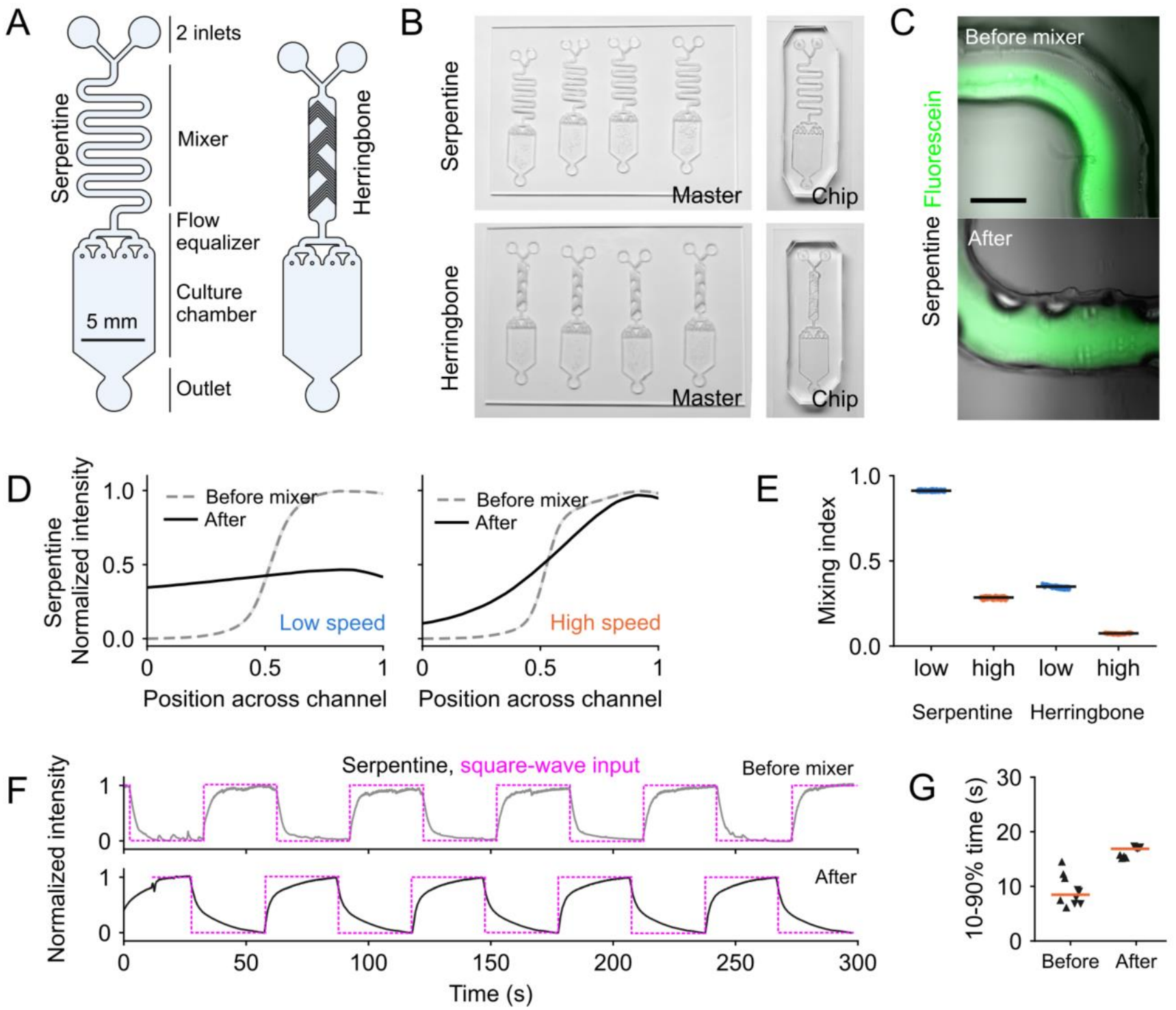
Laser-cut mixers mix or switch two solutions. (A) Designs of the two-inlet device: inlets, mixer (serpentine, left; staggered herringbone, right), flow equalizer, culture chamber and outlet. (B) PMMA masters (four per glass slide) and PDMS devices with the serpentine (top) and her-ringbone (bottom) mixer. (C) Fluorescein (green) overlaid on brightfield before and after the serpentine. Scale bar, 0.5 mm. (D) Fluorescence across the channel before (gray, dashed) and after (black) the serpentine, at 20 µL/min (low speed) and 200 µL/min (high speed); mean of 101 frames (3 s), normalized; position normalized to the analyzed window. (E) Mixing index after each mixer at the two flow rates; points, frames; bars, medians. (F) Fluorescence over time before and after the serpentine when the inputs alternate every 30 s (dotted, magenta, input); the two positions were recorded one after the other. (G) 10–90% time of each transition in F; triangles up, rises; down, falls; bars, medians. One device per design shown, representative of >10 from the same masters. Data in S1 Data.

Before the mixer, the two solutions flowed side by side (Fig 7C and 7D). The serpentine mixed them almost completely at 20 µL/min, but much less at 200 µL/min, when the solutions spend ten times less time in the mixer (Fig 7D and 7E). The herringbone mixed less than the serpentine at both flow rates (Fig 7E). We then asked whether the same device could also switch between the two solutions. We alternated the inputs every 30 s and followed the fluorescence before and after the serpentine (Fig 7F). The output followed the alternation, with transitions that the mixer smoothed but completed within each half-period (Fig 7G). The same device can therefore deliver either a stable mixture of two solutions or alternating pulses of each.

## Discussion

Microfluidic devices for cell culture often need channels and chambers of hundreds of micrometers, yet their fabrication still depends on clean room equipment. We show that a benchtop laser cutter, an oven and a hotplate are enough to make smooth functional devices in a few hours, and we benchmarked each step so that the result can be predicted from the design. The line energy of the laser sets the depth and width of a cut in each master material. PDMS cured just enough to be demolded bonds to glass without any surface treatment when curing is completed in contact with it, and corona activation of the glass raises the bond from about 400 to about 900 mbar. The master material sets the surface of the device, but even the rough walls of tape masters did not perturb laminar flow. With these rules, the same workflow produced devices for compartmentalized co-culture, for laminar flow at defined shear and for mixing and switching of solutions.

Laser-cut and laser-machined masters have been described before [7,10–13,15,16]. In direct machining of PMMA, the width and depth of CO_2_-laser channels have been measured and modeled as functions of power, speed and number of passes [23–25]. For laser-cut masters, earlier calibrations varied laser power and focus at a fixed speed in PMMA [7], varied power and speed one at a time for PMMA molds [14] and engraved masters [37], or measured the track width in tape [10]. A full grid of power and speed shows that the energy per unit length summarizes both settings (S4 Fig), as proposed for PMMA molds [14], and extends this rule to polystyrene and tape. Unlike in direct machining, the cut piece itself becomes the master, so what matters is whether a setting cuts through the material and how much width it removes; we measured both, in rigid sheets and in tape, on a benchtop laser cutter. Partial curing has been used to bond PDMS layers to each other [28], also in bench-top workflows [31]; here, it also bonded PDMS to glass-bottom dishes without any treatment, and activation of the glass alone with a corona device raised the bond to about 900 mbar. Finally, we tested what the calibration alone cannot predict: the surface and the flow of the finished devices.

The main gain is time. With soft lithography, each new design waits for a photomask and a clean-room session; with a laser-cut master, a modified design is cut in minutes, and a bonded device is ready on the same day (Table 1). Other routes also avoid the clean room (compared in Table 1). Resin 3D printers cost no more than a benchtop laser cutter and, unlike laser-cut masters, make molds with several heights and three-dimensional shapes in one print [8]. Most printing resins, however, inhibit the curing of PDMS, so each mold needs a post-treatment that depends on the resin: UV exposure and baking up to 2 days [38], or a paint coating that limits features to about 100 µm [8]. Printed surfaces can also reduce the transparency of PDMS replicas [39]. Photolithography with UV-curable nail gel needs only a UV lamp, a printed photomask, spacers and a solvent wash, and gives features of about 100 µm and heights up to about 1 mm [40], a range similar to that of laser-cut masters, but each new design needs a new photomask. Laser-cut masters need neither a photomask nor a post-treatment, and with PMMA masters the replicas copy surfaces as smooth as glass on both the channel ceiling and the bonding face (Fig 4C).

**Table 1.** Time and cost of a PDMS device made by soft lithography, on 3D-printed or nail-gel molds, or with laser-cut masters. Times and costs of this work are from the Materials and methods (Time and cost estimates). Values for 3D-printed molds are from [8,38,39] and from reseller prices of the Formlabs Form 4 printer, and values for nail-gel photolithography from [40]. Materials per device cost about 3.40 USD for all methods (PDMS and glass-bottom dish). All methods also need an oven, a vacuum desiccator and a hotplate.

|  | **Soft lithography** | **3D-printed molds** | **Nail-gel photolithog-raphy** | **Laser-cut masters (this work)** |
|---|---|---|---|---|
| **Design to master** | 2 days–2 weeks | 2.5 h–2 days | 1–7 days (printed mask) | 0.25–3 h |
| **Master to bonded de-vice** | 2–4 h | About 4 h | About 2 h | About 2.5 h |
| **Equipment cost (USD)** | Mask aligner, >120,000, or clean-room fees; plasma cleaner, 6,000–30,000 | Resin printer, 1,000–5,500 | UV lamp, about 20 | Laser cutter, about 3,000; corona de-vice (optional), about 1,800 |
| **Materials per master (USD)** | 60–180; about 600 with a chrome mask | About 1–2 | Under 1 | About 0.25 |
| **Mold post-treatment** | Silanization, usually | UV and baking, or a coating | Solvent wash | None |
| **Smallest lat-eral feature** | A few µm | About 100 µm | About 100 µm | 0.2 mm (tape); 0.4–0.5 mm (sheets) |
| **Feature height** | 1–100 µm; several heights | Any; several heights and 3D shapes | 80 µm–1 mm; one height | 0.1–2 mm; one per layer, plus partial cuts |
| **Mold surface** | Smooth | Printed layers | Smooth or rough, de-pending on the mask | Glass-smooth faces (PMMA) |

The workflow presented here has a defined range of application. The smallest feature is set by the cut width, about 0.2 mm in tape and 0.4–0.5 mm in sheets, and each master layer has one height. This range covers the chambers, channels, reservoirs and mixers of many culture devices, and partial cuts add a second height, as for the phase guides. Laser settings are specific to each cutter and material, but a single grid of 49 cuts recalibrates the method on another instrument. Tape masters are the fastest to make, because they are cut on their support, and they give the most regular partial cuts; they need cleaning with solvent, their walls are rough, and PVC releases hydrogen chloride when cut [41], so tape should be cut only in an enclosed cutter with exhaust. PMMA masters give smoother surfaces. Partially cured PDMS may contain uncross-linked oligomers that can leach into the medium [42]; the bonding step continues the cure, but we did not measure the remaining fraction. Without a corona device, the replica can also be stamped onto a thin layer of uncured PDMS on the glass before baking, a form of adhesive bonding that has sealed PDMS and glass devices [28,43]; we did not test it. When pressures above 1 bar are needed, oxygen plasma remains the better choice.

Because a master is a vector drawing, designs can be shared and adapted as PDF files using common software. We provide the designs of all devices in this study, as well as other common designs, as an open library of SVG files that can be cut directly or modified (Data and code availability). Photolithography remains necessary for micrometer features. For devices with features of hundreds of micrometers or more, laser-cut masters let biology laboratories make, test and modify their own devices without a specialized fabrication pipeline [9], and quickly adapt the device to the experiment rather than the inverse [8].

## Materials and methods

### Device designs

Devices were drawn as vector graphics in Inkscape and saved as SVG files. Composite shapes were merged (Union) so that only the outer outline was cut, and the files were imported into the laser-cutter software. Design dimensions are nominal, without compensation for the cut width. The same U-shaped test pattern was used for the calibration of PMMA, PS and PVC tape. All design files are available on Figshare (Data and code availability).

### Master materials

Masters were made from 1-mm and 0.5-mm PMMA sheets (Langaelex acrylic sheets), 1-mm and 0.5-mm white polystyrene sheets (Tamiya Pla-Plate) and yellow PVC insulating tape (Cellpack, cat. no. 145787; nominal thickness 150 µm per layer, of which 40 µm adhesive). Glass slides served as supports. For sheets, double-sided adhesive tape (3M 300LSE; about 100 µm) was laminated to one face of the sheet, and its liner was kept until after cutting. For tape masters, the tape was applied to the glass in one layer for the calibration and in three layers (0.40 mm, measured) for the channel replica and the three-channel device, and pressed to remove air bubbles.

### Laser cutting

Masters were cut with a desktop CO_2_ laser cutter (xTool P2S; 55-W glass tube; wavelength 10.64 µm) controlled with xTool Creative Space, with air assist and one pass per line. The distance between the laser head and the material was set with the distance measurement of the instrument. For the calibration, seven powers and seven speeds were combined for each material (49 conditions): 16, 30, 44, 58, 72, 86 and 100% of 55 W and 35 to 245 mm/s in steps of 35 mm/s for PMMA and PS; 1, 7, 14, 21, 28, 35 and 42% and 516 to 600 mm/s in steps of 14 mm/s for PVC tape. The tape grid was cut in Engrave mode and the tape device masters in Cut mode. The nominal line energy was E = P/v, with P the nominal power (percentage × 55 W) and v the translation speed. Masters in 0.5-mm PMMA (leak-test chip, two-inlet devices and three-channel device) were cut at 44% and 35 mm/s. In three layers of tape, the channel replica was cut at 25% and 75 mm/s and the outline of the three-channel device at 2% and 250 mm/s. Each phase guide was two parallel lines 0.2 mm apart, cut partway into the master at 30% and 100 mm/s in PMMA (Cut mode) and at 100% and 350 mm/s in tape (Engrave mode).

### Assembly of masters

After cutting, PMMA or PS features were released from the sheet, the liner was removed, and the features were pressed onto a clean glass support along their full length. For tape masters, the tape outside the features was removed with fine forceps, and the remaining features were pressed onto the glass. Tape masters were cleaned with isopropanol to remove adhesive residues before casting (S5 Fig). Masters were inspected under a stereomicroscope, and masters with incomplete cuts, visible melting or delamination were discarded.

### Scanning electron microscopy

PMMA and PS sheets were broken across the cuts with a razor blade to expose cross-sections. PVC tape was applied at the edge of a PMMA slide and cut across that edge, so that each cross-section lay at the edge. Masters and casts of the three-channel device were imaged from above, and casts were cut along and across the phase guides with a steel razor. Samples were coated with 10 nm of palladium and imaged on a Zeiss Merlin SEM (EPFL CIME) at 2 kV with the HE-SE2 detector. Cross-sections of laser cuts were imaged at 113× (2.42 µm/pixel; PMMA and PS) and 400× (0.684 µm/pixel, two images at 475×; PVC tape), and tilted views at 20° at 97–100× (2.73–2.82 µm/pixel; PMMA and PS) or 200× (1.37 µm/pixel; PVC tape). Masters and casts of the three-channel device were imaged at 0° or 20° tilt and 57–70× (3.9–4.8 µm/pixel), and sections at 75–400× (0.68–3.64 µm/pixel). Overlapping images of the sections were stitched in Fiji [44].

### Cut measurements

W, H and the width at the base were measured in ImageJ (version 1.54p) on one cut per condition, calibrated with the scale bar of each image: H from the top surface to the apex of the groove, W at the level of the top uncut surface, and the width at the base for cuts through the sheet. Cuts through the 1-mm sheet (H = 1000 µm) were treated as censored at the sheet thickness, and tape cuts with H > 110 µm as cuts through the PVC film. For S1–S3 Fig, cross-sections were aligned on the top surface, centered on the cut, resampled to one pixel size per material and shown with one linear contrast per material.

### Surface roughness and phase-guide height

Roughness was measured on tilted SEM views of masters and casts (3–4 images per condition, from two imaging sessions). Regions were drawn on flat surfaces, on cut edges and on glass (the glass support next to masters, and PDMS molded against glass next to casts). In each region, the SEM signal was band-pass filtered with Gaussian filters (σ = 4 and 20 µm; features of about 10–60 µm), and roughness was the standard deviation of the band-passed signal divided by the mean SEM signal of the region. The median of the regions of one image gave one value per image. The channel height under the phase guide was manually measured along the stitched longitudinal sections of the casts at every pixel. The two ends of each trace, outside the phase guide, were excluded.

### PDMS casting and demolding

PDMS (Sylgard 184, Dow) base and curing agent were mixed at 10:1 by mass and degassed under vacuum until no bubbles were visible. PDMS was poured over the master to a thickness of about 5 mm and cured in an oven at 70 °C for 50 min (standard protocol, used for all devices in Figs 4–7) or, for the curing-time experiments, for 10 to 120 min in 10-min steps. After cooling, the PDMS was peeled from the master, and ports were made with a coring punch (Rapid-Core, Well-Tech; inner diameter 0.5 mm, outer diameter 0.8 mm).

### PDMS bonding

PDMS replicas were bonded to 35-mm glass-bottom dishes: Ibidi (cat. no. 81218), FluoroDish (WPI, cat. no. FD35-100) and VWR confocal dishes (cat. no. 734-2904). For thermal bonding, partially cured replicas were placed in contact with the dish and heated at 85 °C for 45 min on a hotplate (Stuart SB160), either without treatment or after activation of the dish surface for 45 s with a handheld corona device (MTP3, STTS) held about 5 mm from the surface; the PDMS was not treated. For oxygen-plasma bonding, the PDMS and the glass of Ibidi dishes were treated in a plasma cleaner (Diener electronic Femto, type SLS) at 100 W and 1 mbar for 45 s, brought into contact immediately and heated at 85 °C for 45 min on the hotplate.

### Leak pressure

Leak pressure was measured on chips with one straight channel (11.8 mm long and 2.0 mm wide in the design; 0.58 mm high), one inlet and one outlet. Stainless-steel 18G pin connectors were inserted into the ports. The inlet was connected to a pressure controller (LineUP Flow EZ, Fluigent; maximum 1000 mbar), and the outlet to an open reservoir. The channel was filled with 0.1 mM fluorescein in water. Pressure was increased in 50-mbar steps every minute, and the step at which liquid leaked from the device was recorded as the leak pressure. Devices that held 1000 mbar were recorded as ≥1000 mbar. Oxygen-plasma-bonded devices were tested in the same way up to 2.3 bar with a pressure generator (FLPG plus, Fluigent). Three devices were tested per condition.

### Time and cost estimates

Drawing a design took 5 min to 1 h, depending on its complexity, and cutting the master and mounting it on glass about 10 min (at most about 70 min). Mixing the PDMS took 5 min, degassing on the master 40 min, curing 50 min, peeling and punching 5 min, corona activation 45 s and bonding on the hotplate 45 min, about 2.5 h per device from the master. The materials cost about 0.20 CHF per master (0.10 CHF for the glass support and 0.10 CHF for the plastic sheet; the double-sided tape is negligible) and 2.60 CHF per device (2.20 CHF for a 35-mm glass-bottom dish and 0.60 CHF for 2.7 g of PDMS).

### Three-channel device: cell culture and imaging

A549 human lung adenocarcinoma cells (ATCC, CCL-185) were transduced with a lentiviral vector encoding mCherry to obtain stable fluorescent reporter expression. Cells were maintained in F-12K medium (Kaighn’s modification of Ham’s F-12; Gibco, Thermo Fisher Scientific) supplemented with 2 mM L-glutamine, 10% (v/v) fetal bovine serum (FBS), 100 U/mL penicillin and 100 µg/mL streptomycin (all Gibco). MRC-5 human lung fibroblasts (ECACC, 05072101; Merck) were cultured in DMEM with GlutaMAX supplemented with 10% (v/v) FBS, 100 U/mL penicillin and 100 µg/mL streptomycin (all Gibco). For seeding, A549 cells (10 × 10^6^ cells/mL) and MRC-5 cells (2.5 × 10^6^ cells/mL) were resuspended in a gel of 80% (v/v) Matrigel (Growth Factor Reduced, phenol red-free, LDEV-free; Corning, 356231) and 20% (v/v) rat tail collagen I (Corning) conjugated to silicon rhodamine (SiR; Spirochrome), kept on ice and loaded into the two lateral channels of the device, one after the other. The device was incubated at 37 °C and 5% CO_2_ for 1 h to let the gels polymerize. The inlet and outlet of the central channel were then fitted with stainless-steel pins and silicone tubing (inner diameter 0.8 mm; Ibidi, 10841), connected to a 10-mL glass bottle with a double-connector cap holding 10 mL of complete F-12K medium, which served as reservoir and bubble trap. The central channel was perfused at 50 µL/min with a peristaltic pump (HPM100, Innofluid) inside the incubator for at least 3 days. Devices were imaged on a Nikon Ti2 microscope with an X-Light V3 spinning-disk unit and a Plan Apo λD 20×/0.8 objective, in brightfield and in the Cy5 (excitation 640 nm, emission 684 nm; SiR-collagen) and Cy3 (excitation 545 nm, emission 595 nm; mCherry) channels.

### Image analysis of the three-channel device

The stitched images of the device before and 12h after the start of perfusion (0.65 µm/pixel, averaged over 5 × 5 pixels) were aligned on the collagen channel by rotation and template matching, and cropped to the same 11.05-mm square. The collagen signal was smoothed (Gaussian, σ = 3 pixels) and scaled to 0 in the PDMS and 1 at the median of the gel (Otsu threshold). In each image row, the gel front on each side was the innermost pixel above 0.5, and the gap was the distance between the two fronts. The phase-guide section comprised the rows in which the gap was below 2.36 mm in both images, halfway between its design values at the phase guides (2.13 mm) and between the channel walls (2.60 mm). The collagen profile across the device was averaged over this section. For display, each channel was scaled linearly between its 1st and 99.5th percentiles; no other processing was applied.

### Channel replica and particle image velocimetry

The channel replica (3.8 × 21 mm) was cut in three layers of PVC tape, cast in PDMS and bonded to a 24 × 60 mm glass coverslip. The commercial channel was one channel of an Ibidi µ-Slide VI 0.4 (cat. no. 80600). Water containing 1-µm fluorescent microspheres (Fluoresbrite, Polysciences, cat. no. 17154-10; 1:10,000) was pumped at 25, 50 and 100 µL/min with a syringe pump (WPI, AL8000). Images were acquired on a Nikon Eclipse Ti2 microscope with a Plan Fluor 4×/0.13 objective and a Kinetix camera (Teledyne Photometrics) at 6.5 µm/pixel, with frame intervals of 3.5 ms (replica) and 2.3 ms (commercial slide). Velocity fields were computed with OpenPIV (version 0.26.1; doi:10.5281/zenodo.3566451) from frame pairs two frames apart, with a 16-pixel interrogation window, a 19-pixel search area and a 9-pixel overlap, after a high-pass filter (σ = 5 pixels); vectors with a signal-to-noise ratio below 1.8 were replaced by the local mean. All recordings were processed with the same parameters, and every second field was used, so that no frame entered two fields (34–52 fields per recording). The analysis region was 0.3–4.2 mm across and 0–0.8 mm along the channel.

The streamwise velocity was averaged over time and along the flow to give the profile across the channel. The plateau velocity was the mean of the profile over the central 2 mm, and the width of the profile was taken where it crossed half the plateau. Profiles were compared with the series solution for laminar flow in a rectangular channel, averaged over the channel depth and over the 104-µm interrogation window. Vorticity, ω = ∂v/∂x − ∂u/∂y, was computed by central differences after averaging the velocity over 3 × 3 vectors and normalized as ωW/Ū, with W = 3.8 mm and Ū = Q/(wh) the mean velocity expected from the flow rate Q and the nominal cross-section. Velocity fluctuations were the root-mean-square deviation from the time average in the central 2 mm, divided by Ū. The PIV noise was estimated by reprocessing the raw recordings at 25 µL/min at two frame separations (2 and 8 frames), assuming that the displacement error does not depend on the separation. The variation of the wall vorticity along the flow was computed in the grid column of maximal time-averaged vorticity, after subtraction of the variance expected from the standard error of the time average. Wall shear rates were the maximal slopes of the profile near each wall, with 95% confidence intervals from 2,000 resamplings of the PIV fields. The floor shear stress was estimated as 6µQ/(wh^2^), and the Reynolds number as ρŪD_h_/µ, with D_h_ = 2wh/(w + h) and the density and viscosity of water at 20 °C.

### Two-inlet devices and mixing index

The two-inlet devices had channels about 0.6 mm high. The serpentine mixer was 0.35 mm wide, and the herringbone mixer 2.8 mm wide, with grooves 0.12 mm deep in the channel ceiling. One inlet carried 0.1 mM fluorescein sodium and the other distilled water. For steady mixing, flow was driven at 20 and 200 µL/min with pressure controllers (LineUP Flow EZ, Fluigent). Fluorescence was imaged before and after the mixer on a Nikon Ti2 microscope with a CFI Plan Apo λD 10×/0.45 objective and a Kinetix camera, as series of 101 frames at 30-ms intervals, and intensity profiles across the channel were extracted in Fiji (1.54p). Each profile was cropped to pixels 100– 600, smoothed with a 15-pixel moving average and normalized as (I − I_min_)/(I_max_ − I_min_), with I_min_ and I_max_ the 2nd and 98th percentiles of the time-averaged profile before the mixer at the same flow rate. The mixing index of each frame after the mixer was MI = 1 − σ_after_/σ_before_, where σ is the standard deviation of the normalized profile across the channel and σ_before_ its mean over the frames before the mixer. MI = 0 means no mixing and MI = 1 a uniform profile. For the herringbone device, one series before the mixer, recorded at the low flow rate, served for both flow rates. Illumination was not flat-field corrected.

For the alternating input, two syringe pumps (Aladdin AL-8000, World Precision Instruments), one per inlet, were switched every 30 s by a Python script (Figshare) and images were recorded every 34–72 ms. The mean intensity in a region of interest was measured in Fiji, normalized between the 2nd and 98th percentiles of each time course and smoothed over 9 frames, and the 10–90% time of each transition that reached both plateaus was measured.

### Statistics

Analyses were made in Python 3.13 with NumPy, pandas, SciPy, scikit-image, OpenPIV and Matplotlib (versions in requirements.txt), with one Jupyter notebook per figure (Figshare). The replicate unit is given in each legend: laser conditions for the calibration, devices for leak pressure, images for roughness, and recordings for PIV and mixing. We used three rank-based tests, which make no assumption on the distribution and suit small samples: Spearman’s rank correlation for trends, the two-sided Mann–Whitney U test for two groups, and the Kruskal–Wallis test for more than two groups. Leak pressures of devices that held the maximum pressure were ranked at that value. Depth and width were fitted as linear functions of line energy by least squares, excluding cuts through the material and conditions without a measurable cut. PIV and mixing data come from one recording per condition and are reported without tests. Resampling used a fixed random seed.

## Author contributions

JMGA designed the experimental plan, wrote the manuscript and contributed to the experiments; JAM performed most of the experiments and edited the draft; CA prepared the 3-channel device experiment. All authors co-designed the three channel device.

## Acknowledgments

We thank the ISREC at EPFL for funding for the laser cutter equipment, and Elisa Oricchio for support and discussions. We thank Pierre Gönczy and his team for feedback support throughout this work and for hosting. We thank Guillermina Rochelle Ramirez-San-Juan and Benedetta Noferi for help setting up the laser cutter experiments and sourcing materials. We thank Aikaterini Maria Kourkoulou for setting up the PIV video recording. The SEM experiments have been conducted on the equipment in the Electron Microscopy Center of EPFL, CIME. We thank CIME for support and assistance, particularly Fabienne Bobard. We would like to thank the EPFL BioImaging & Optics Core Facility and in particular Angélique Vinit and José Artacho for help and assistance using the spinning disk confocal. This work was funded by the SNF Ambizione program PZ00-3_233294 and by the BNF qualification program from the university of Bern.

## Data and code availability

The data underlying all figures are in S1 Data. Raw images, the analysis notebooks and the vector design files of all devices (SVG) are available on Figshare (https://doi.org/10.6084/m9.figshare.34944154).

## Use of AI tools

We used Claude (Anthropic Claude Opus 5.5, October 2026) to write and debug the Python code in the analysis notebooks (image processing, measurement extraction, statistics and plotting), to edit the wording, and to search the scientific literature. The authors defined all analyses, edited the code and rerun each notebook from the raw data. No AI tool was used to generate or modify primary data or images. The authors take full responsibility for the content.

## Supporting information

**S1 Fig.**
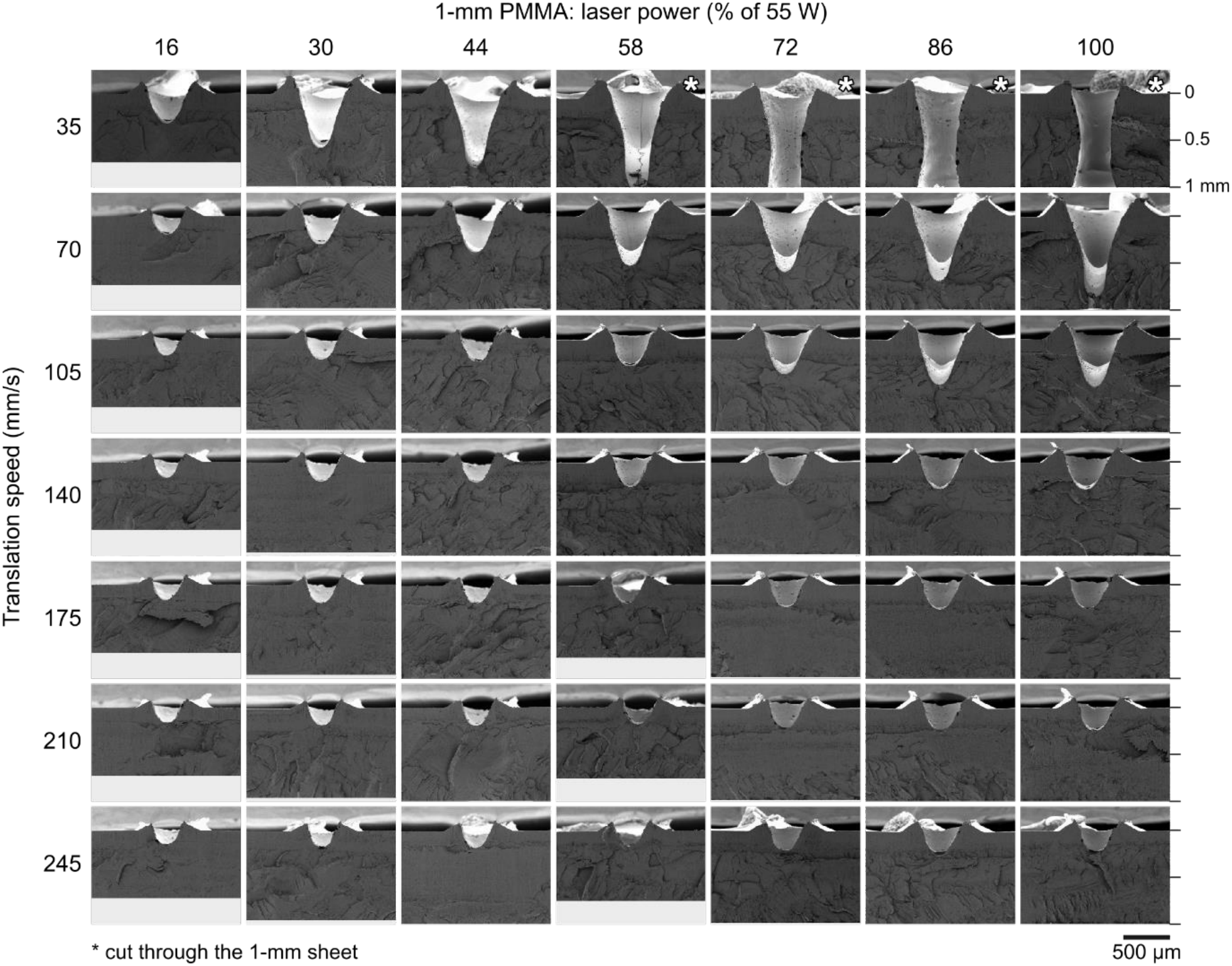
SEM cross-sections of laser cuts in 1-mm PMMA over the full power–speed grid. One cut per condition; laser power in columns (% of 55 W), translation speed in rows (mm/s). Each tile is aligned on the top surface of the sheet and centered on the cut; ticks on the right mark the surface and 0.5 and 1.0 mm below it. One linear contrast for all tiles; light gray, outside the imaged field. Asterisks, cuts through the sheet. Scale bar, 500 µm. Measured values in Fig 2D.

**S2 Fig.**
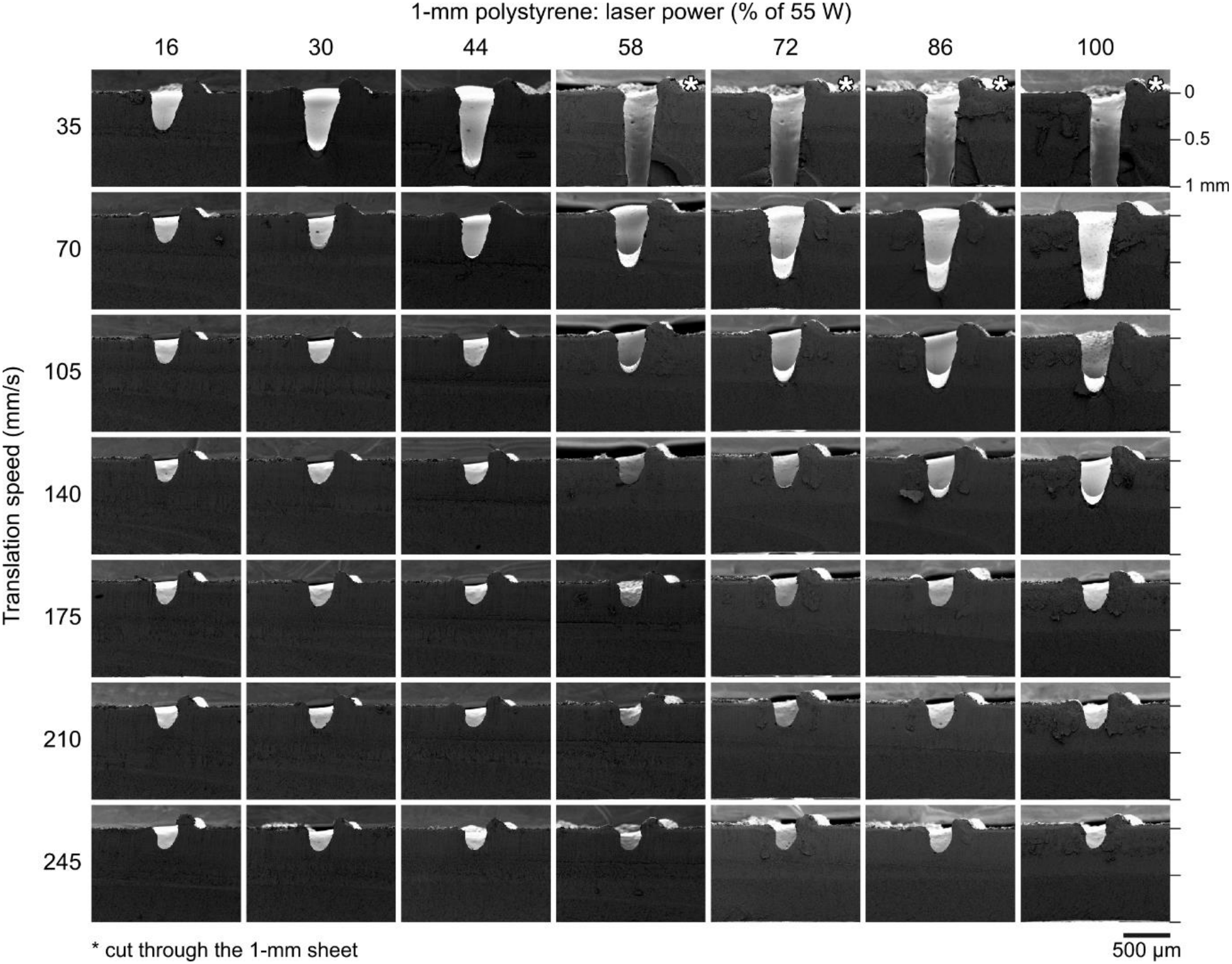
SEM cross-sections of laser cuts in 1-mm polystyrene over the full power–speed grid. As S1 Fig. Measured values in Fig 2F.

**S3 Fig.**
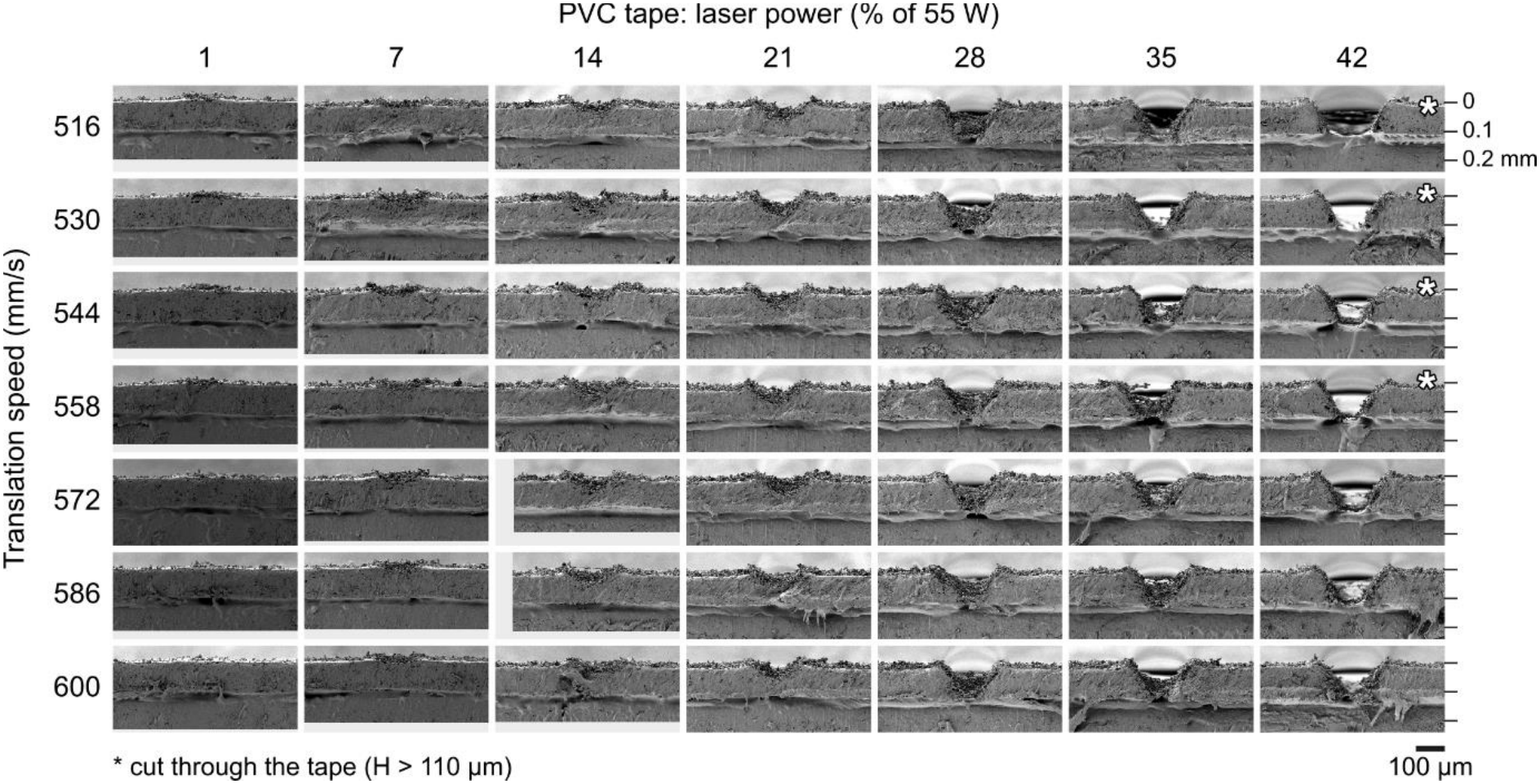
SEM cross-sections of laser cuts in PVC tape over the full power–speed grid. As S1 Fig; ticks mark the surface and 0.1 and 0.2 mm below it, and asterisks cuts through the PVC film (H > 110 µm). Images at 475× were resampled to the pixel size of the 400× images. Scale bar, 100 µm. Measured values in Fig 2E.

**S4 Fig.**
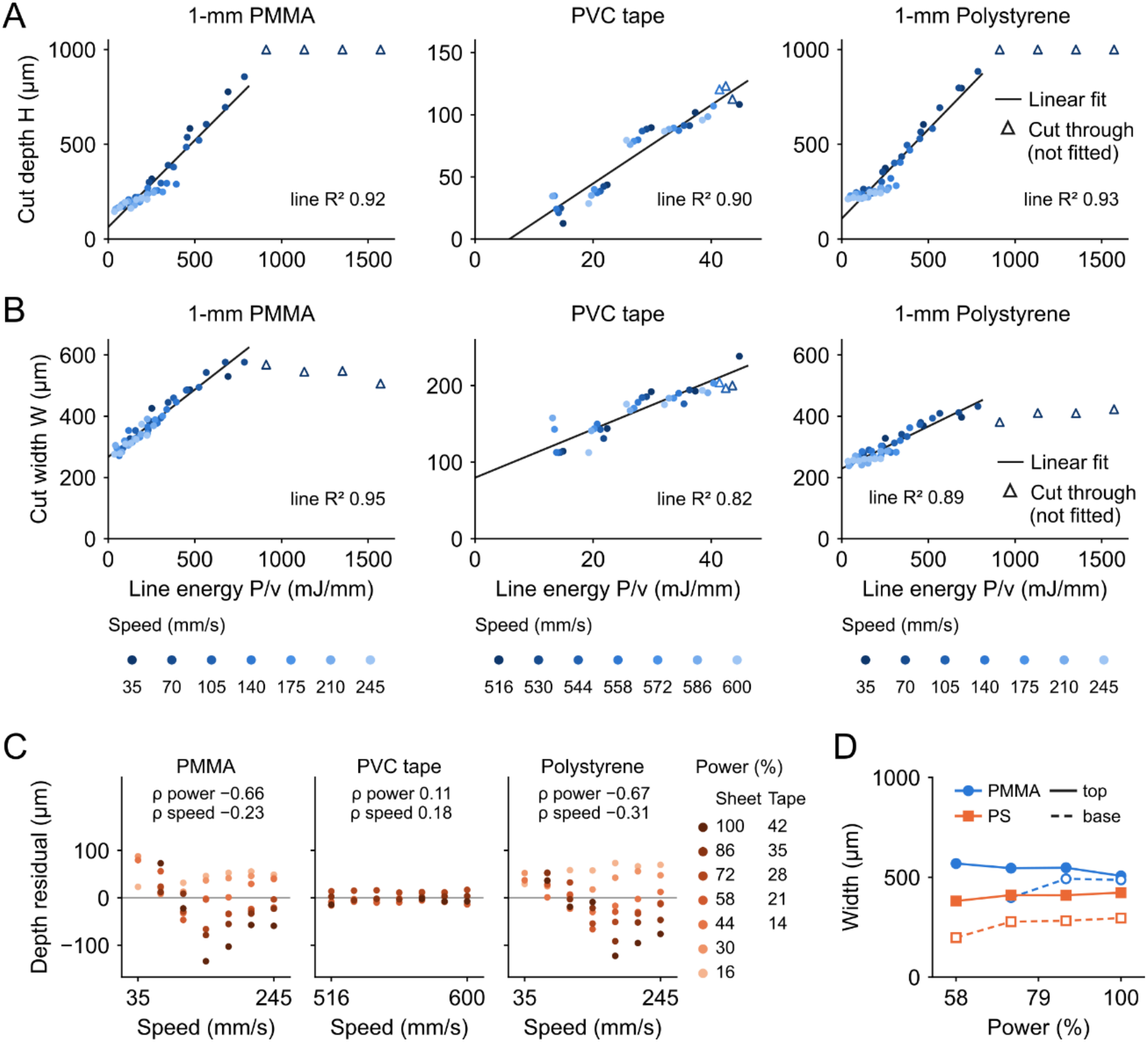
Cut depth and width increase linearly with line energy. (A) Cut depth H and (B) cut width W as a function of the nominal line energy P/v for PMMA, PVC tape and PS (color, translation speed). Solid lines, straight-line fits (R^2^ = 0.90–0.93 for depth and 0.82–0.95 for width); open triangles, cuts through the sheet or the tape, excluded from all fits. (C) Residuals of the depth fits in A as a function of speed; color, laser power. ρ, Spearman correlation of the residuals with power and with speed (n = 45 conditions for PMMA and PS, 31 for PVC tape); in the sheets, depth fell below the line at high power. (D) Width at the top surface (filled symbols, solid lines) and at the base (open symbols, dashed lines) of cuts through 1-mm PMMA and PS at 35 mm/s. One cut per condition. Data in S1 Data.

**S5 Fig.**
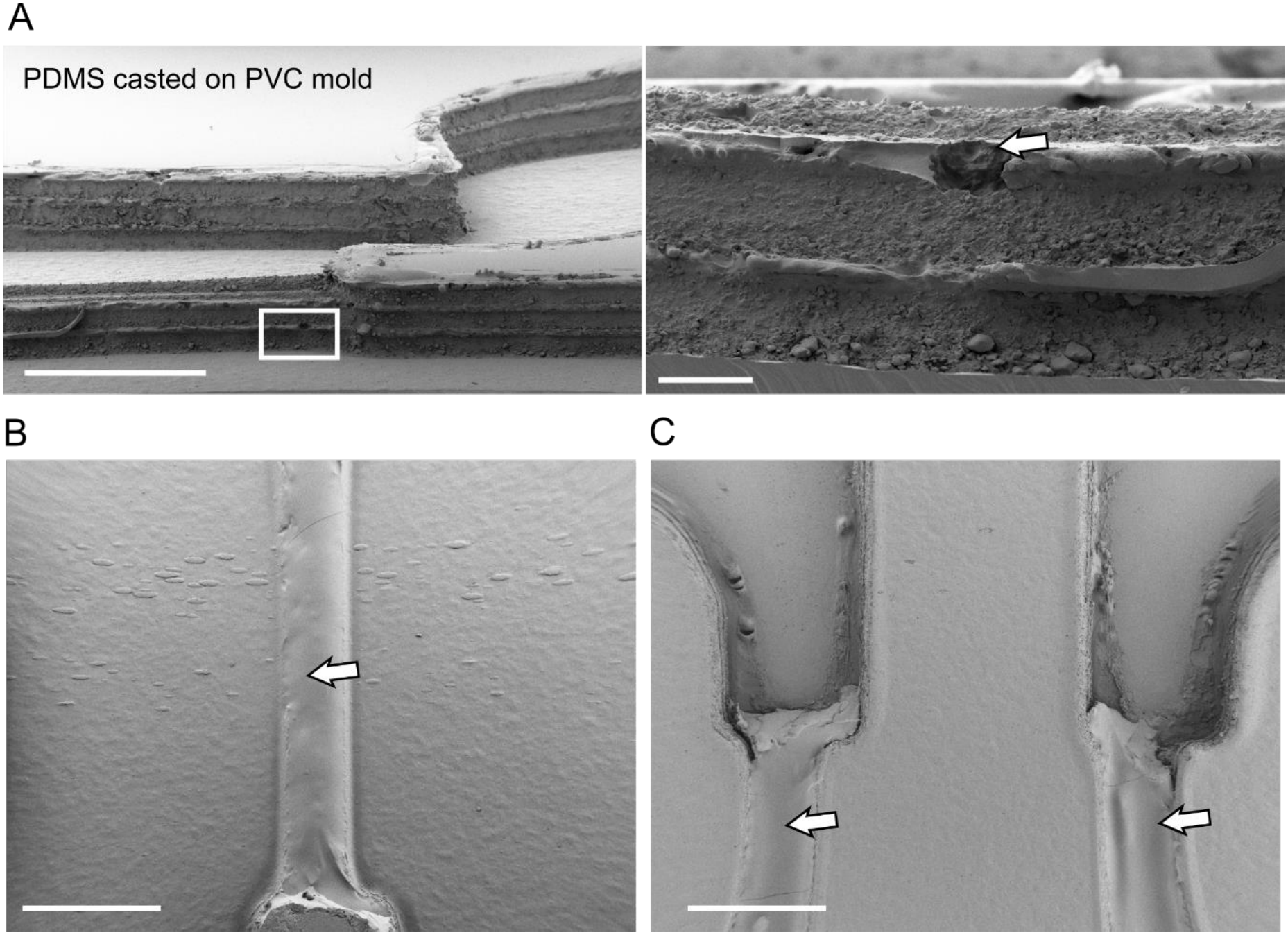
Defects of multilayer PVC tape masters. (A) Longitudinal section of a PDMS cast from a three-layer PVC tape master (15° tilt; left) and enlargement of the boxed region (right): the cast reproduces the layered edge of the tape stack; arrow, fracture in the PDMS. (B) PDMS cast and (C) PVC tape master at the phase guides, when the master was not cleaned with solvent; arrows, adhesive left on the master (C) and reproduced in the cast (B). Scale bars, 1 mm (A left, B and C) and 100 µm (A right).

**S1 Data. Underlying data for all quantitative figure panels.** Excel workbook with one sheet per figure panel.

## References

1. Whitesides GM. The origins and the future of microfluidics. Nature. 2006;442: 368–373. doi:10.1038/nature05058

2. Sackmann EK, Fulton AL, Beebe DJ. The present and future role of microfluidics in biomedical research. Nature. 2014;507: 181–189. doi:10.1038/nature13118

3. McDonald JC, Whitesides GM. Poly(dimethylsiloxane) as a Material for Fabricating Microfluidic Devices. Acc Chem Res. 2002;35: 491–499. doi:10.1021/ar010110q

4. Xia Y, Whitesides GM. Soft Lithography. Angewandte Chemie International Edition. 1998;37: 550–575. doi:10.1002/(SICI)1521-3773(19980316)37:5<550::AID-ANIE550>3.0.CO;2-G

5. Duffy DC, McDonald JC, Schueller OJA, Whitesides GM. Rapid Prototyping of Microfluidic Systems in Poly(dimethylsiloxane). Anal Chem. 1998;70: 4974–4984. doi:10.1021/ac980656z

6. Ren K, Zhou J, Wu H. Materials for Microfluidic Chip Fabrication. Acc Chem Res. 2013;46: 2396–2406. doi:10.1021/ar300314s

7. Matellan C, Del Río Hernández AE. Cost-effective rapid prototyping and assembly of poly(methyl methacrylate) microfluidic devices. Sci Rep. 2018;8: 6971. doi:10.1038/s41598-018-25202-4

8. Hagemann C, Bailey MCD, Carraro E, Stankevich KS, Lionello VM, Khokhar N, et al. Low-cost, versatile, and highly reproducible microfabrication pipeline to generate 3D-printed customised cell culture devices with complex designs. Khosla C, editor. PLoS Biol. 2024;22: e3002503. doi:10.1371/journal.pbio.3002503

9. Walsh DI, Kong DS, Murthy SK, Carr PA. Enabling Microfluidics: from Clean Rooms to Makerspaces. Trends in Biotechnology. 2017;35: 383–392. doi:10.1016/j.tibtech.2017.01.001

10. Luo LW, Teo CY, Ong WL, Tang KC, Cheow LF, Yobas L. Rapid prototyping of microfluidic systems using a laser-patterned tape. J Micromech Microeng. 2007;17: N107–N111. doi:10.1088/0960-1317/17/12/N02

11. Qin S, Ou G, Wang B, Li Z, Hu R, Li Y, et al. Photolithography-free fabrication of photoresist-mold for rapid prototyping of microfluidic PDMS devices. Chinese Chemical Letters. 2022;33: 987–989. doi:10.1016/j.cclet.2021.07.045

12. Samuel R, Thacker CM, Maricq AV, Gale BK. Simple and cost-effective fabrication of microvalve arrays in PDMS using laser cut molds with application to *C. elegans* manipulation in microfluidics. J Micromech Microeng. 2014;24: 105007. doi:10.1088/0960-1317/24/10/105007

13. Isiksacan Z, Guler MT, Aydogdu B, Bilican I, Elbuken C. Rapid fabrication of microfluidic PDMS devices from reusable PDMS molds using laser ablation. J Micromech Microeng. 2016;26: 035008. doi:10.1088/0960-1317/26/3/035008

14. Guler MT, Inal M, Bilican I. CO2 laser machining for microfluidics mold fabrication from PMMA with applications on viscoelastic focusing, electrospun nanofiber production, and droplet generation. Journal of Industrial and Engineering Chemistry. 2021;98: 340–349. doi:10.1016/j.jiec.2021.03.033

15. Joanni E, Peressinotto J, Domingues PS, Setti GDO, Jesus DPD. Fabrication of molds for PDMS microfluidic devices by laser swelling of PMMA. RSC Adv. 2015;5: 25089–25096. doi:10.1039/C5RA03122B

16. Gerber LC, Kim H, Riedel-Kruse IH. Microfluidic assembly kit based on laser-cut building blocks for education and fast prototyping. Biomicrofluidics. 2015;9: 064105. doi:10.1063/1.4935593

17. Jiang J, Zhan J, Yue W, Yang M, Yi C, Li C-W. A single low-cost microfabrication approach for polymethylmethacrylate, polystyrene, polycarbonate and polysulfone based microdevices. RSC Adv. 2015;5: 36036–36043. doi:10.1039/C5RA02220G

18. Bartholomeusz DA, Boutte RW, Andrade JD. Xurography: rapid prototyping of microstructures using a cutting plotter. J Microelectromech Syst. 2005;14: 1364–1374. doi:10.1109/JMEMS.2005.859087

19. Comina G, Suska A, Filippini D. PDMS lab-on-a-chip fabrication using 3D printed templates. Lab Chip. 2014;14: 424–430. doi:10.1039/C3LC50956G

20. Faustino V, Catarino SO, Lima R, Minas G. Biomedical microfluidic devices by using low-cost fabrication techniques: A review. Journal of Biomechanics. 2016;49: 2280–2292. doi:10.1016/j.jbiomech.2015.11.031

21. Nguyen H-T, Thach H, Roy E, Huynh K, Perrault CM-T. Low-Cost, Accessible Fabrication Methods for Microfluidics Research in Low-Resource Settings. Micromachines. 2018;9: 461. doi:10.3390/mi9090461

22. Rodríguez CF, Guzmán-Sastoque P, Gantiva-Diaz M, Gómez SC, Quezada V, Muñoz-Ca-margo C, et al. Low-cost inertial microfluidic device for microparticle separation: A laser-Ablated PMMA lab-on-a-chip approach without a cleanroom. HardwareX. 2023;16: e00493. doi:10.1016/j.ohx.2023.e00493

23. Klank H, Kutter JP, Geschke O. CO2-laser micromachining and back-end processing for rapid production of PMMA-based microfluidic systems. Lab Chip. 2002;2: 242. doi:10.1039/b206409j

24. Snakenborg D, Klank H, Kutter JP. Microstructure fabrication with a CO_2_ laser system. J Micromech Microeng. 2004;14: 182–189. doi:10.1088/0960-1317/14/2/003

25. Yuan D, Das S. Experimental and theoretical analysis of direct-write laser micromachining of polymethyl methacrylate by CO2 laser ablation. Journal of Applied Physics. 2007;101: 024901. doi:10.1063/1.2409621

26. Bhattacharya S, Datta A, Berg JM, Gangopadhyay S. Studies on surface wettability of poly(dimethyl) siloxane (PDMS) and glass under oxygen-plasma treatment and correlation with bond strength. J Microelectromech Syst. 2005;14: 590–597. doi:10.1109/JMEMS.2005.844746

27. Haubert K, Drier T, Beebe D. PDMS bonding by means of a portable, low-cost corona system. Lab Chip. 2006;6: 1548. doi:10.1039/b610567j

28. Eddings MA, Johnson MA, Gale BK. Determining the optimal PDMS–PDMS bonding technique for microfluidic devices. J Micromech Microeng. 2008;18: 067001. doi:10.1088/0960-1317/18/6/067001

29. Haddadi E, Moradi M, Karimzad Ghavidel A, Karimzad Ghavidel A, Meiabadi S. Experimental and parametric evaluation of cut quality characteristics in CO2 laser cutting of polystyrene. Optik. 2019;184: 103–114. doi:10.1016/j.ijleo.2019.03.040

30. Mushtaq RT, Wang Y, Rehman M, Khan AM, Mia M. State-Of-The-Art and Trends in CO2 Laser Cutting of Polymeric Materials—A Review. Materials. 2020;13: 3839. doi:10.3390/ma13173839

31. Saem S, Zhu Y, Luu H, Moran-Mirabal J. Bench-Top Fabrication of an All-PDMS Microfluidic Electrochemical Cell Sensor Integrating Micro/Nanostructured Electrodes. Sensors. 2017;17: 732. doi:10.3390/s17040732

32. Vulto P, Podszun S, Meyer P, Hermann C, Manz A, Urban GA. Phaseguides: a paradigm shift in microfluidic priming and emptying. Lab Chip. 2011;11: 1596. doi:10.1039/c0lc00643b

33. Trietsch SJ, Israëls GD, Joore J, Hankemeier T, Vulto P. Microfluidic titer plate for stratified 3D cell culture. Lab Chip. 2013;13: 3548–3554. doi:10.1039/c3lc50210d

34. Santiago JG, Wereley ST, Meinhart CD, Beebe DJ, Adrian RJ. A particle image velocimetry system for microfluidics. Experiments in Fluids. 1998;25: 316–319. doi:10.1007/s003480050235

35. Hersen P, McClean MN, Mahadevan L, Ramanathan S. Signal processing by the HOG MAP kinase pathway. Proc Natl Acad Sci USA. 2008;105: 7165–7170. doi:10.1073/pnas.0710770105

36. Stroock AD, Dertinger SKW, Ajdari A, Mezić I, Stone HA, Whitesides GM. Chaotic Mixer for Microchannels. Science. 2002;295: 647–651. doi:10.1126/science.1066238

37. Nguyen QT, Pham UT, Nguyen DT, Tran NT, Dang LD, Hoang HM, et al. Fabrication of micro-droplet generating microfluidic master mold using reverse laser engraving. Microsyst Technol. 2026;32: 26. doi:10.1007/s00542-025-05975-1

38. Venzac B, Deng S, Mahmoud Z, Lenferink A, Costa A, Bray F, et al. PDMS Curing Inhibition on 3D-Printed Molds: Why? Also, How to Avoid It. Anal Chem. 2021;93: 7180–7187. doi:10.1021/acs.analchem.0c04944

39. Parthiban P, Vijayan S, Doyle PS, Hashimoto M. Evaluation of 3D-printed molds for fabrication of non-planar microchannels. Biomicrofluidics. 2021;15: 024111. doi:10.1063/5.0047497

40. Nguyen H-T, Thach H, Roy E, Huynh K, Perrault CM-T. Low-Cost, Accessible Fabrication Methods for Microfluidics Research in Low-Resource Settings. Micromachines. 2018;9: 461. doi:10.3390/mi9090461

41. Kiefer M, Moss CE. Laser generated air contaminants released during laser cutting of fabrics and polymers. J Laser Appl. 1997;9: 7–13. doi:10.2351/1.4745440

42. Regehr KJ, Domenech M, Koepsel JT, Carver KC, Ellison-Zelski SJ, Murphy WL, et al. Biological implications of polydimethylsiloxane-based microfluidic cell culture. Lab Chip. 2009;9: 2132. doi:10.1039/b903043c

43. Wu H, Huang B, Zare RN. Construction of microfluidic chips using polydimethylsiloxane for adhesive bonding. Lab Chip. 2005;5: 1393–1398. doi:10.1039/b510494g

44. Preibisch S, Saalfeld S, Tomancak P. Globally optimal stitching of tiled 3D microscopic image acquisitions. Bioinformatics. 2009;25: 1463–1465. doi:10.1093/bioinformatics/btp184

